# Versatile open software to quantify cardiomyocyte and cardiac muscle contraction *in vitro* and *in vivo*

**DOI:** 10.1101/160754

**Authors:** L. Sala, B.J. van Meer, L.G.J. Tertoolen, J. Bakkers, M. Bellin, R.P. Davis, C. Denning, M.A.E. Dieben, T. Eschenhagen, E. Giacomelli, C. Grandela, A. Hansen, E.R. Holman, M.R. M. Jongbloed, S.M. Kamel, C.D. Koopman, Q. Lachaud, I. Mannhardt, M.P.H. Mol, V.V. Orlova, R. Passier, M.C. Ribeiro, U. Saleem, G.L. Smith, C.L. Mummery, F.L. Burton

## Abstract

Contraction of muscle reflects its physiological state. Methods to quantify contraction are often complex, expensive and tailored to specific models or recording conditions, or require specialist knowledge for data extraction. Here we describe an automated, open-source software tool (MUSCLEMOTION) adaptable for use with standard laboratory and clinical imaging equipment that enables quantitative analysis of normal cardiac contraction, disease phenotypes and pharmacological responses. MUSCLEMOTION allowed rapid and easy measurement of contractility in (i) single cardiomyocytes from primary adult heart and human pluripotent stem cells, (ii) multicellular 2D-cardiomyocyte cultures, 3D engineered heart tissues and cardiac organoids/microtissues *in vitro* and (iii) intact hearts of zebrafish and humans *in vivo*. Good correlation was found with conventional measures of contraction in each system. Thus, using a single method for processing video recordings, we obtained reliable pharmacological data and measures of cardiac disease phenotype in experimental cell- and animal models and human echocardiograms.

## Introduction

The salient feature of cardiomyocytes (CMs) is their ability to undergo cyclic contraction and relaxation, a feature critical for cardiac function. In many research laboratories and clinical settings it is therefore essential that cardiac contraction can be quantified at multiple levels, from single cells to multicellular or intact cardiac tissues. Measurement of contractility is relevant for analysis of disease phenotypes, cardiac safety pharmacology, and longitudinal measures of cardiac function over time, both *in vitro* and *in vivo*. In addition, human genotype-phenotype correlations, investigation of cardiac disease mechanisms and the assessment of cardiotoxicity are increasingly performed on human induced pluripotent stem cells (hiPSCs) derived from patients^1-3^. Many of these studies are carried out in non-specialist laboratories so that it is important that analysis methods are simplified such that they can be used anywhere with access to just standard imaging equipment. Here, we describe a single method with high versatility that can be applied to most imaging outputs of cardiac contraction likely to be encountered in the laboratory or clinic.

Electrical and calcium signals are usually quantified *in vitro* using established technologies such as patch clamp electrophysiology, multi electrode arrays, cation-sensitive dyes or cation-sensitive genetic reporters^4^. Although experimental details differ among laboratories, the values for these parameters are with some approximations comparable across laboratories, cardiomyocyte source and cell culture configuration (e.g. single cells, multicellular 2-Dimensional (2D) CM monolayers, 3-Dimensional (3D) cultures)^5^,^6^. However, there is no comparable method for measuring cardiac contraction across multiple platforms, despite this being a crucial functional parameter affected by many diseases or drugs^7^. We have developed a method to address this that is built on existing algorithms and is fully automated, but most importantly can be used on videos, image stacks or image sequences loaded in the open source image processing program ImageJ^8^. Moreover, it is an open source, dynamic platform that can be expanded, improved and integrated for customized applications. The method, called MUSCLEMOTION, determines dynamic changes in pixel intensity between image frames and expresses the output as a relative measure of displacement during muscle contraction and relaxation. We applied the concept to a range of biomedical- and pharmacologically relevant experimental models that included single hPSC-CMs, patterned- or 2D cultures of hPSC-CMs, cardiac organoids, engineered heart tissues (EHTs) and isolated adult rabbit CMs. Results were validated by comparing outputs of the tool with those from three established methods for measuring contraction: optical flow, post deflection and fractional shortening of sarcomere length. These methods have been tailored to (or only work on) specific cell configurations. Traction force microscopy, fractional shortening of sarcomere length and microposts are predominantly suitable for single cells^8^,^9^. Cardiomyocyte edge or perimeter detection is suitable for adult CMs but challenging for immature hPSC-CMs due to poorly defined plasma membrane borders and concentric contraction^10^, while large post deflection is suitable for EHTs or small cardiac bundles^11^ but less so for single cells. Our MUSCLEMOTION software by contrast can be used for all of these applications without significant adaptions. Furthermore, it can be used for multi-parameter recording conditions and experimental settings using transmitted light microscopy, fluorescent membrane labeling, fluorescent beads embedded in soft substrates or patch clamp video recordings. Drug responses to positive and negative inotropic agents were evaluated across four different laboratories in multiple cell configurations using MUSCLEMOTION with reliable predictions of drug effects from all laboratories. Furthermore, MUSCLEMOTION was also applicable to optical recordings of zebrafish hearts *in vivo*, where it represented a significant time-saving in analysis, and in human echocardiograms. This versatile tool thus provides a rapid and straightforward way to detect disease phenotypes and pharmacological responses *in vitro* and *in vivo.*

## Methods

Extended methods are in the Supplementary Information. The datasets generated and/or analyzed during the current study are available from the corresponding authors on reasonable request.

### Code Availability

MUSCLEMOTION source code is included in the Supplementary Material and is available for use and further development.

### Model Cell

The *in silico* cardiomyocyte-like model (**Fig. 1d,f,g**) was created using Blender v2.77.

**Figure 1.**
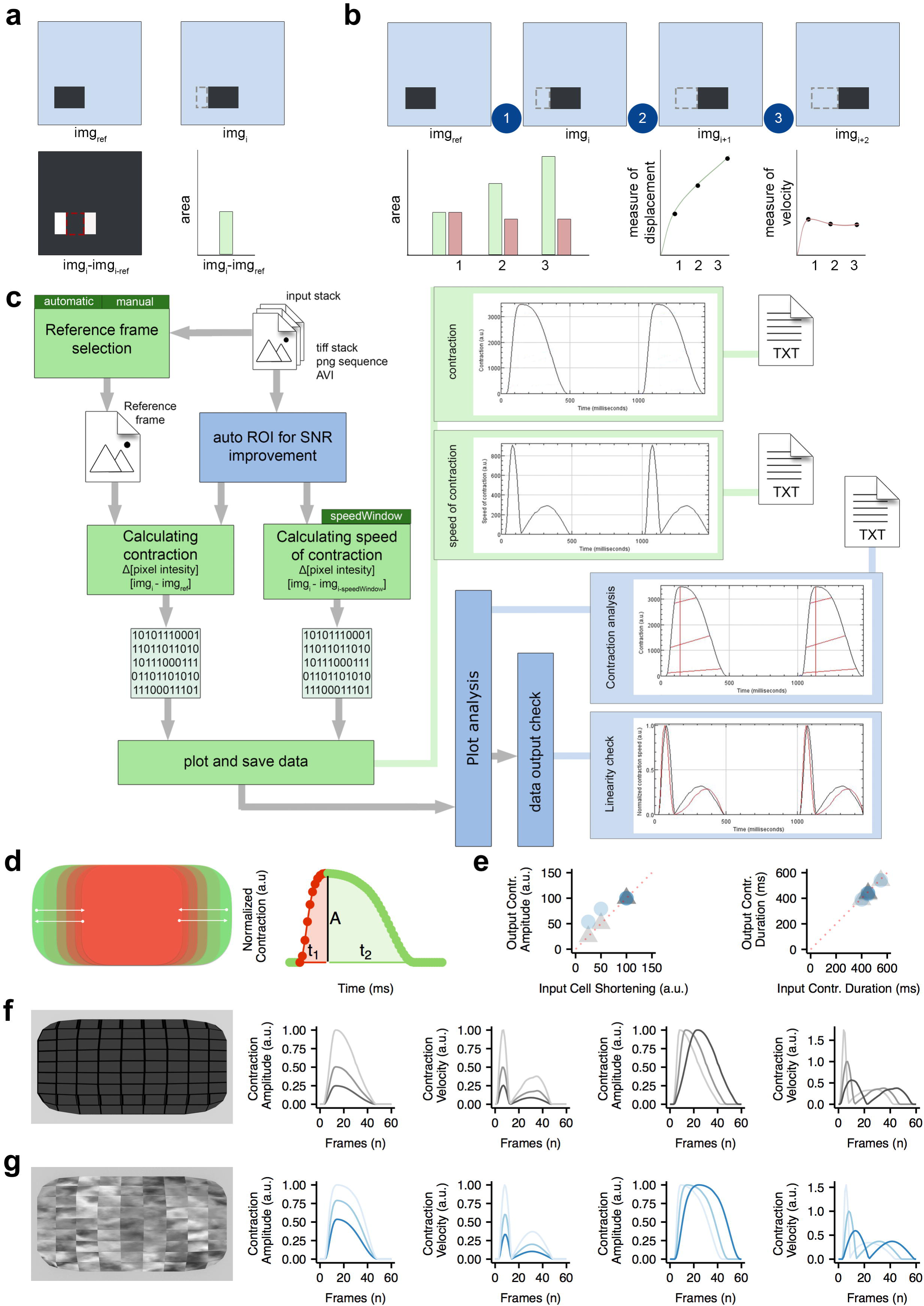
Algorithm construction and validation. **a)** Principle of pixel intensity difference by subtraction of *img*_*ref*_ of *img*_*i*_ and measurement of the non-zero area after image subtraction. **b)** Principle of using pixel intensity difference as a measure of displacement and as a measure of velocity. **c)** Schematic overview of MUSCLEMOTION. Green blocks indicate basic steps of the algorithm. Dark green blocks indicate important user input choices. Plots within light green blocks indicate results. Optional steps are shown in blue blocks, with graphical representation of the analysed parameters indicated by red lines. Three result files are generated containing the raw data: “contraction.txt”, “speed-of-contraction.txt” and “overview-results.txt”. Furthermore, three images showing relevant traces and a log file are generated and saved (not shown in schematic). **d)** Schematic of the contractile pattern of the artificial cell and relative parameters corresponding to amplitude of contraction (A), time-to-peak (t_1_) and relaxation time (t_2_). **e)** Correlation between input (x axis) and output (y axis) parameters used to validate MUSCLEMOTION with two artificial cells. **f-g)** Frame representing the two artificial cells built for MUSCLEMOTION validation and their relative output parameters.

### Optical Flow analysis

Optical flow analysis was implemented in LabVIEW as described by Hayakawa et al.^12^,^13^.

### hPSC Culture and Differentiation

hPSCs from multiple independent cell lines (**Table S1**) were differentiated to CMs as previously described^14-17^, or with the Pluricyte^®^ Cardiomyocyte Differentiation Kit (Pluriomics b.v.) according to the manufacturer’s protocol. Experiments were performed at 18-30 days after initiation of differentiation, depending on the cell source and configuration. Pluricytes^®^ were kindly provided by Pluriomics b.v.

### Patch Clamp Recordings on hPSC-CMs

Electrophysiological recordings of isolated hPSC-CMs were performed as previously described^16^.

### Movement of embedded beads

Gelatin-patterned polyacrylamide gels containing fluorescent beads were generated and analyzed as described previously^18^.

### Monolayers of hPSC-CMs

25k-40k cells were plated per Matrigel-coated glass ø10 mm coverslip.

### Cardiac Organoids

Cardiac organoids composed of hPSC-CMs and hPSC-derived endothelial cells, were generated as previously described^17^.

### Adult cardiomyocytes

CMs were isolated from New Zealand White male rabbits as previously described ^19^.

### Membrane labelling

hPSC-CMs were plated on Matrigel-coated glass-bottom 24-well plates and labelled with CellMask Deep Red according to the manufacturer’s instructions.

### Engineered heart tissues

EHTs were generated and analyzed as previously described^14^.

### Zebrafish hearts

Zebrafishes hearts were recorded, treated and analysed as previously described ^23^.

### Echocardiograms

Anonymized ultrasounds of 5 adult patients were selected from the echocardiography database of the Leiden University Medical Center.

### Statistics

One-way ANOVA for paired or unpaired measurements was applied to test the differences in means on normalized drug effects. P-values obtained from two-tailed pairwise comparisons were corrected for multiple testing using Bonferroni’s method. Statistical analyses were performed with R v3.3.3. P-values lower than 0.05 were considered statistically significant and indicated with an asterisk (*).

## Results

### Algorithm development

The principle underlying the algorithm of MUSCLEMOTION is the assessment of contraction using an intuitive approach quantifying absolute changes in pixel intensity between a reference frame and the frame of interest, which can be described as

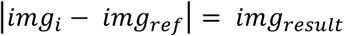

where *img*_*i*_ is the frame of interest, *img*_*ref*_ is the reference frame and _*result*_ is the resulting image. For every pixel in the frame, each reference pixel is subtracted from the corresponding pixel of interest and the difference is presented in absolute numbers. Unchanged pixels result in low (black) values, while pixels that are highly changed result in high (white) values (**Fig. 1a**). Next, the mean pixel intensity of the resulting image is measured. This is a quantitative measure of how much the pixels have moved compared to the reference frame: more white pixels indicate more changing pixels and, thus, more displacement. When a series of images is analysed relative to the same reference image, the output describes the accumulated displacement over time (measure of displacement, **Fig. 1b**).

However, if a series of images is analysed with a reference frame that depends on the frame of interest (e.g. *img*_*ref*_*= img*_*i-1*_), this results in a measure of the relative displacement per interframe interval. We defined this parameter as contraction velocity (measure of velocity, **Fig. 1b**).

Since velocity is the first derivative of displacement in time, the first derivative of the measure of displacement should resemble the measure of velocity derived from image calculations. To test the linearity of the method, three movies of moving blocks were analysed. The block moved back and forth at two different speeds in each direction (where *v*_*2*_*= 2* · *v*_1_): i) along the x-axis, ii) along the y-axis and iii) along both axes (**Movie S1**). As expected, the measure of displacement and velocity showed a linear correlation (**Fig. S1**). This does not hold when the position of the block in *img*_*i*_ does not overlap the position of the block in *img*_*ref*_, with a consequent saturation in the measure of displacement (i.e. max pixel white value, **Fig. S2**). Therefore, comparison of the differentially derived velocities should approximately overlap in the absence of pixel saturation. This was used as a qualitative parameter to determine whether the algorithm outputs were reliable.

### Algorithm implementation

MUSCLEMOTION was then modified to handle typical experimental recordings by (i) improving the signal-to-noise ratio (SNR), (ii) automating reference frame selection and (iii) programming built-in checks to validate the generated output data (**Fig. 1c**). The SNR was increased by isolating the pixels of interest in a three-step process: i) maximum projection of pixel intensity in the complete displacement stack, ii) creation of a binary image of this maximum projection with a threshold level equal to the mean grey value plus standard deviation and iii) multiplication of the pixel values in this image by the original displacement and speed of the displacement image stack (**Fig. S3**). This process allowed the algorithm to work on a region of interest with movement above the noise level only.

Next, a method was developed to identify the correct *img*_*ref*_ from the speed of displacement image stack by comparing values obtained from the frame-to-frame calculation with their direct neighbouring values, while also checking for the lowest absolute value (**Fig. S4**).

The reliability of MUSCLEMOTION for structures with complex movements was validated using a custom-made contracting 3D “synthetic CM” model (**Fig. 1d,f,g**) that was adapted to produce contractions with known amplitude and duration. Linearity was preserved during the analysis of the contraction and velocity; other output parameters of the analysis matched the input parameters (**Fig. 1e**). A second 3D model (**Fig. 1g**), with a repetitive pattern aimed to create out-of-bounds problems was also generated. As expected, contraction amplitude information here was not linear (**Fig. 1e**), although contraction velocity and temporal parameters did remain linear (**Fig. 1e,g**). To mitigate this problem, we implemented an option for a 10-sigma Gaussian blur filter that can be applied on demand to biological samples that presented highly repetitive patterns (e.g. sarcomeres in adult CMs).

### Algorithm application to multiple cell configurations and correlation with existing gold standards

This set of experiments aimed to investigate the versatility of MUSCLEMOTION and examine how its performance compared with standard measures used in each system: i) optical flow for isolated hPSC-CMs, monolayers and organoids; ii) post deflection for EHT; iii) sarcomere length fractional shortening for adult CMs. Remarkably, standard methods currently used measure only contraction or contraction velocity. Linearity was preserved in all cases during the analyses, demonstrating the reliability of the results (**Fig. S5**).

First, single hPSC-CMs (**Fig. 2a, Movie S2**) exhibited concentric contraction (**Fig. 2a ii**) and contraction velocity amplitudes correlated well with the amplitudes obtained by optical flow analysis (R^2^ = 0.916) (**Fig. 2a v**). In contrast to single cells, the area of displacement for hPSC-CM monolayers was distributed heterogeneously throughout the whole field (**Fig. 2b ii, Movie S3**). Optical flow analysis was compared with our measure of velocity (**Fig. 2b iv**); this showed a good linear correlation (R^2^ = 0.803) (**Fig. 2b v**). Complex (mixed, multicellular) 3D configurations were also investigated by analyzing hPSC-derived cardiac organoids^17^ (**Movie S4**) and EHTs^14^ (**Movie S5**). Cardiac organoids showed moderate levels of displacement throughout the tissue (**Fig. 2c ii**), while the EHTs showed high deflection throughout the bundle (**Fig. 2d ii**). The contraction velocity of the organoids correlated well with the output of optical flow analysis (R^2^ = 0.747, **Fig. 2c v**). Similarly, contraction amplitudes in EHTs showed high linear correlation (R^2^ = 0.819) with the absolute force values derived from measurement of pole deflection (**Fig. 2d v**). Finally, single adult rabbit ventricular CMs were analyzed (**Fig. 2e, Movie S6**). Large displacements were evident around the long edges of the CM (**Fig. 2e ii**). These cells were analyzed with a 10-sigma Gaussian blur filter, which also minimized (unwanted) effects of transverse movements on contraction patterns. Linearity was preserved (**Fig. S5**) despite the repetitive pattern of the sarcomeres and this resulted in accurate measures of both contraction (**Fig. 2e iii**) and speed of contraction (**Fig. 2e iv**). The contraction amplitude of the adult CMs stimulated at 1 Hz correlated well with the output of sarcomeric shortening using fast Fourier transform analysis^20^ (R^2^ = 0.871, **Fig. 2e v**). Thus, the MUSCLEMOTION algorithm yielded data in these initial studies comparable with methods of analysis tailored for the individual platforms.

**Figure 2.**
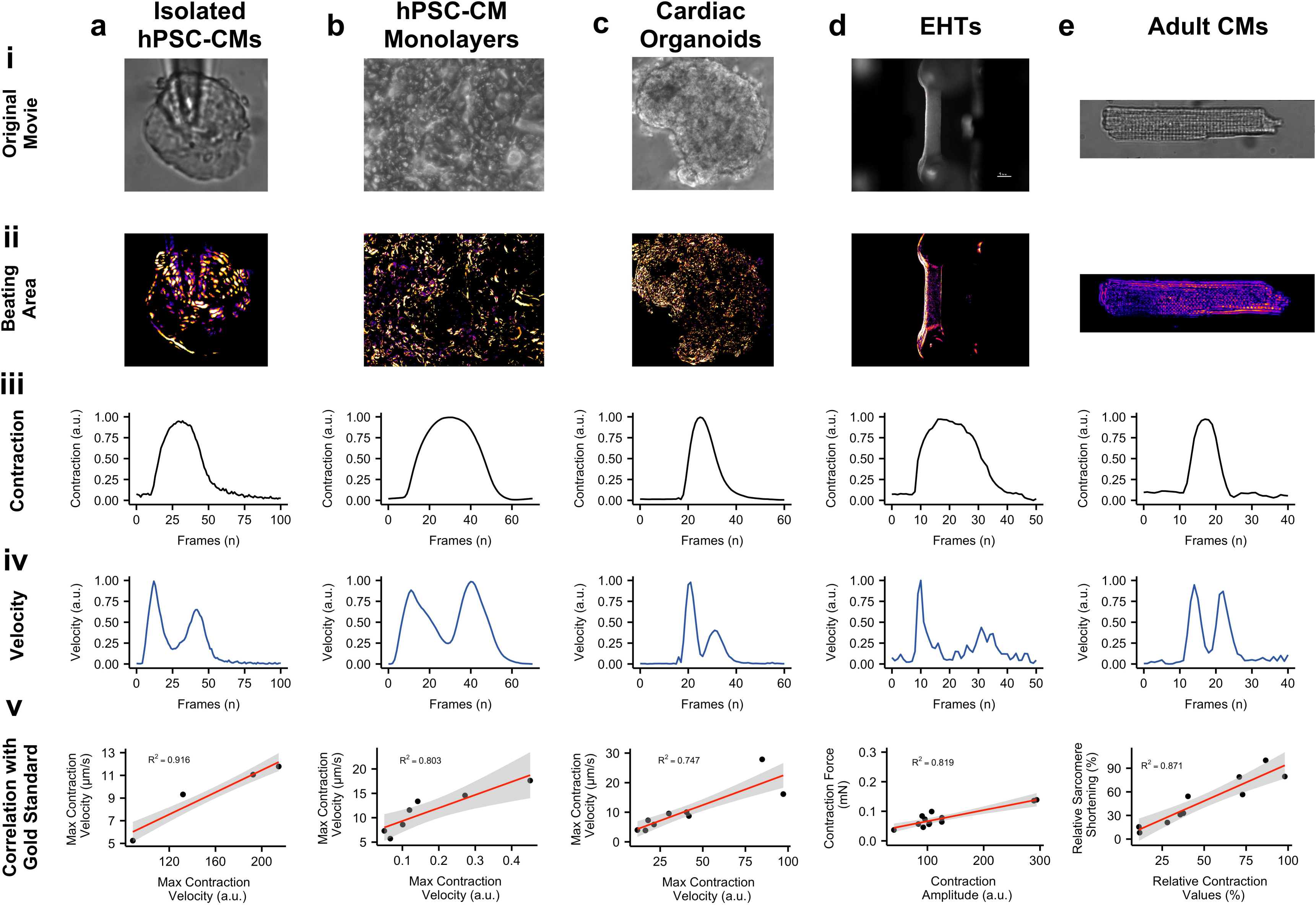
Correlation of results with gold standards. **a)** Brightfield image of isolated hPSC-CMs **(i)**, with maximum projection step visually enhanced with a fire Look Up Table **(ii)**, contraction **(iii)** and velocity **(iv)** profiles of each individual beat have been generated by MUSCLEMOTION and temporally aligned; linear regression analysis between MUSCLEMOTION results (x-axis) and optical flow results (y-axis) **(v)**. **b)** Phase contrast image of hPSC-CM monolayers **(i)**, with maximum projection step visually enhanced with a fire Look Up Table **(ii)**, contraction **(iii)** and velocity **(iv)** profiles of each individual beat have been generated by MUSCLEMOTION and temporally aligned; linear regression analysis between MUSCLEMOTION results (x-axis) and those obtained with optical flow results (y-axis) **(v)**. **c)** Phase contrast image of cardiac organoids **(i)**, with maximum projection step visually enhanced with a fire Look Up Table **(ii)**, contraction **(iii)** and velocity **(iv)** profiles of each individual beat have been generated by MUSCLEMOTION and temporally aligned; linear regression analysis between MUSCLEMOTION results (x-axis) and those obtained with optical flow results (y-axis) **(v)**. **d)** Live view of an EHT during contraction analysis. Scale bar = 1 mm. **(i)**, with maximum projection step visually enhanced with a fire Look Up Table **(ii)**, contraction **(iii)** and velocity **(iv)** profiles of each individual beat have been generated by MUSCLEMOTION and temporally aligned; linear regression analysis between MUSCLEMOTION results (x-axis) and those obtained with post deflection (y-axis) **(v)**. **e)** Brightfield image of adult rabbit CMs **(i)**, with maximum projection step visually enhanced with a fire Look Up Table **(ii)**; contraction **(iii)** and velocity **(iv)** profiles of each individual beat have been generated by MUSCLEMOTION and temporally aligned; linear regression analysis between MUSCLEMOTION results (x-axis) and those obtained from sarcomere fractional shortening calculation with Fast Fourier Transform (y-axis) **(v)**. For details on cell sources and cell lines please refer to the Supplementary Table 1.

### Application of MUSCLEMOTION to multiple imaging and recording platforms

To examine whether MUSCLEMOTION could potentially be used in applications that measure other aspects of CMs functionality in parallel, we first determined the electrophysiological properties of hPSC-CMs using patch clamp whilst recording their contractile properties through video imaging. This allowed simultaneous quantitative measurement of action potentials (APs) and contraction (**Fig. 3a**), for in-depth investigation of their interdependence. We observed a typical^21^ profile of AP followed by its delayed contraction.

**Figure 3.**
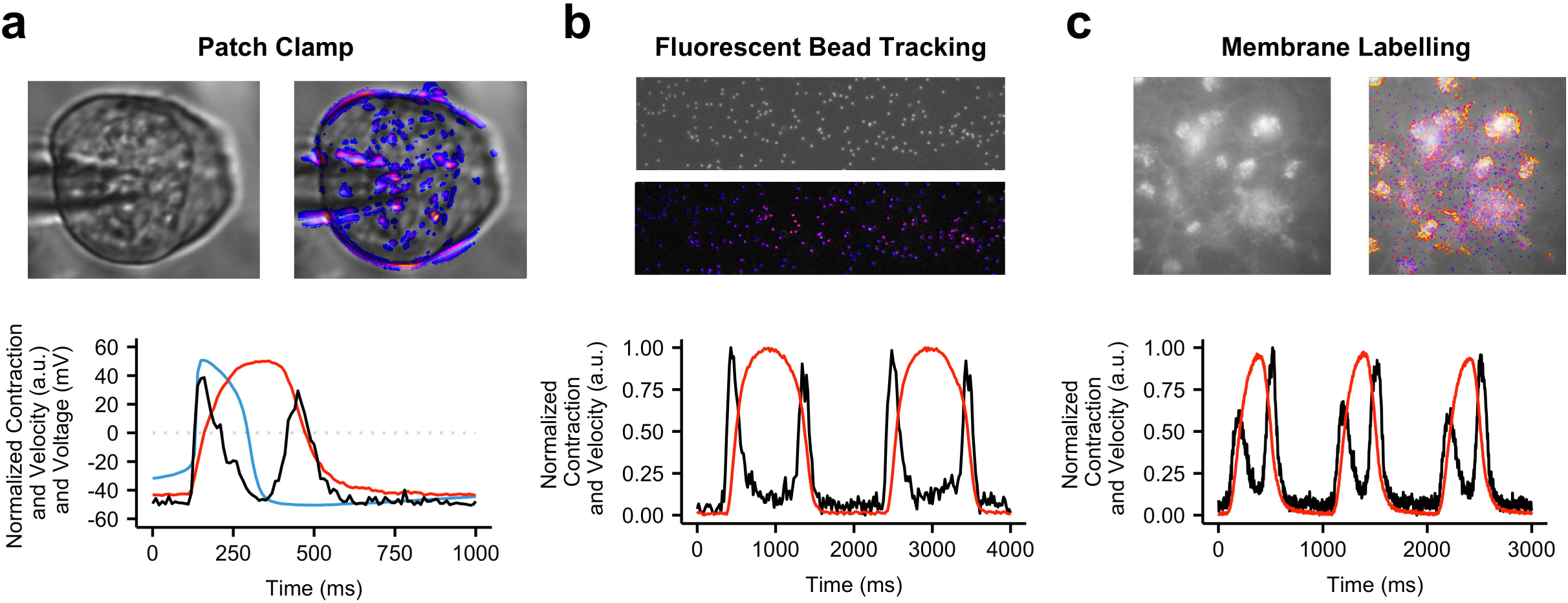
Application of contraction tool to multiple biological situations. Representative examples with enhancement of moving pixels **(top)** and profiles **(bottom)** of contraction **(a-c, red)**, velocity **(a-c, black)** and voltage **(a, blue)** respectively obtained from high speed movies of patched hPSC-CMs **(a)**, aligned hPSC-CMs on polyacrylamide gels with fluorescent beads **(b)** and hPSC-CMs whose membranes have been labelled with CellMask Deep Red **(c)**.For details on cell sources and cell lines please refer to the Supplementary Table 1.

To measure contractile force in combination with contractile velocity in single CMs, we integrated fluorescent beads into polyacrylamide substrates patterned with gelatin (**Fig. 3b**), where the displacement of the beads is a measure of CM contractile force^18^ (**Movie S7**).

Similarly, effective quantification of contraction profiles was obtained for fluorescently labeled hPSC-CM monolayer cultures (**Fig. 3c, Movie S8**), allowing MUSCLEMOTION to be integrated on high speed fluorescent microscope systems for automated data analysis.

### Application of MUSCLEMOTION to drug responses in different cell models in different laboratories

Having shown that MUSCLEMOTION was fit-for-purpose in analyzing contraction over a variety of platforms, we next sought to demonstrate its ability to detect the effects of positive and negative inotropes. This is essential for ensuring the scalability of the tool over multiple platforms, particularly in the context of hiPSC-CMs where regulatory authorities and pharmaceutical companies are interested in using these cells as human heart models for drug discovery, target validation or safety pharmacology^22^. For isoprenaline (ISO) and nifedipine (NIFE) the main parameters of interest are: contraction amplitude (ISO, NIFE), relaxation time (ISO) and contraction duration (NIFE).

The relaxation time of spontaneously beating isolated hPSC-CMs on gelatin patterned polyacrylamide substrates treated with ISO significantly decreased as expected at doses higher than 1 nM. Similar to what has been reported^27^, contraction amplitude decreased at doses higher than 1 nM. NIFE treatment decreased both contraction amplitude and duration starting from 3 nM, respectively (**Fig. 4a**). In paced (1.5 Hz) hPSC-CMs monolayers, no significant effects were measured after addition of ISO on either relaxation time or contraction amplitude. NIFE caused a progressive decrease in contraction duration and amplitude in a concentration-dependent manner starting at 100 nM (**Fig. 4b**). Similarly, cardiac organoids paced at 1.5 Hz showed no significant effects on both relaxation time and contraction amplitude with ISO, while both parameters decreased after NIFE, starting from 100 nM and 300 nM, respectively (**Fig. 4c**). EHTs paced at 1.5 times baseline frequency and analyzed with MUSCLEMOTION showed a positive inotropic effect starting from 1 nM ISO and a negative inotropic effect starting at 30 nM NIFE as previously reported^14^ (**Fig. 4d**).

**Figure 4.**
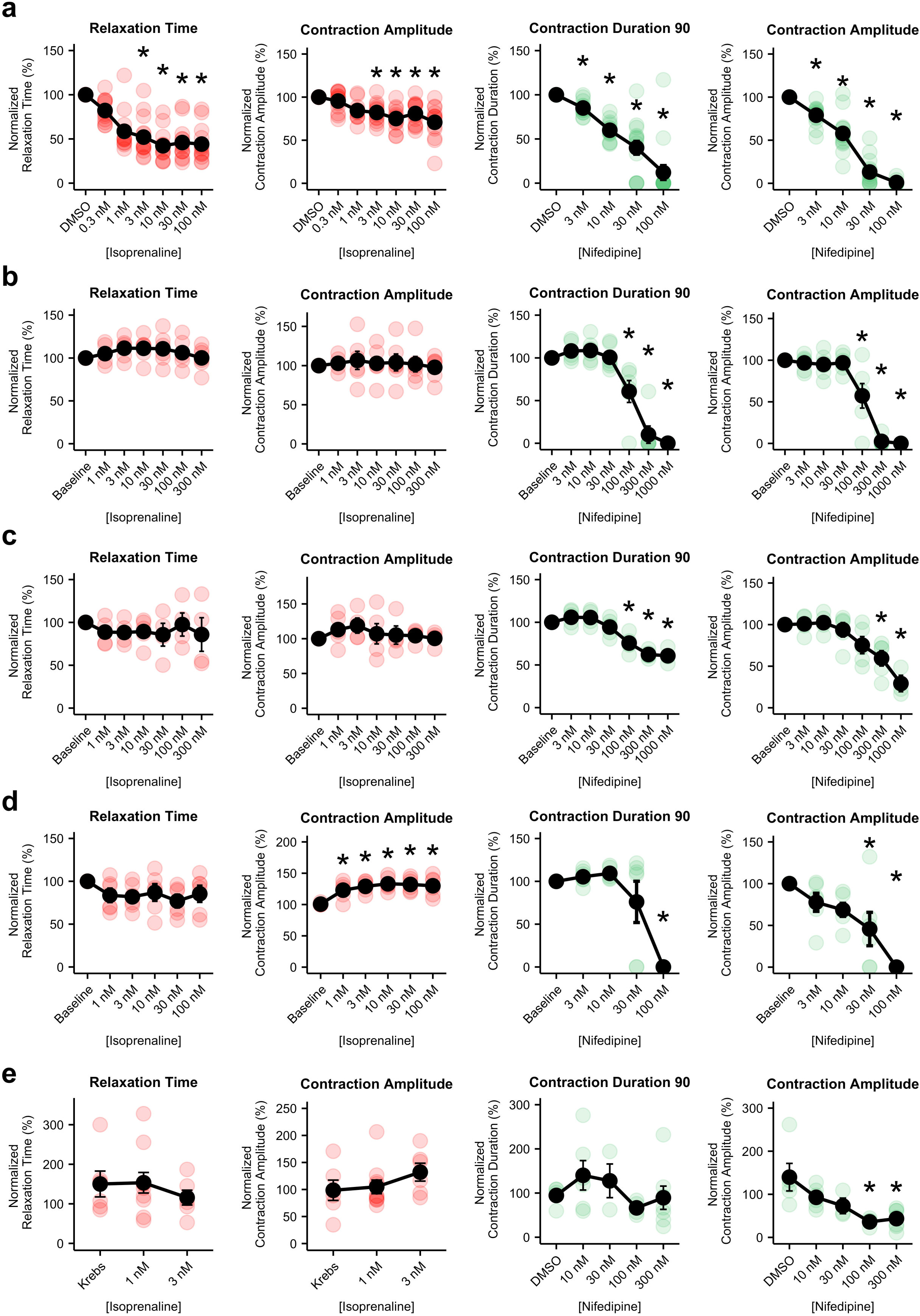
Pharmacological challenge with positive and negative inotropic compounds. **a)** Average dose-response curves (**black traces**) and single measurements for several parameters obtained in isolated, spontaneously beating, aligned hPSC-CMs treated with isoprenaline **(left, red)** and nifedipine **(right, green)**. **b)** Average dose-response curves (**black traces**) and single measurements for several parameters obtained from monolayers of hPSC-CMs treated with isoprenaline **(left, red)** and nifedipine **(right, green)**. **c)** Average dose-response curves (**black traces**) and single measurements for several parameters obtained in cardiac organoids treated with isoprenaline **(left, red)** and nifedipine **(right, green)**. **d)** Average dose-response curves (**black traces**) and single measurements for several parameters obtained in EHTs treated with isoprenaline **(left, red)** and nifedipine **(right, green)**. **e)** Average dose-response curves (**black traces**) and single measurements for several parameters obtained in adult rabbit CMs treated with isoprenaline **(left, red)** and verapamil **(right, green)**. Average data points (**black**) represent mean ± standard error of mean. For details on cell sources and cell lines please refer to the Supplementary Table 1. Data information: P-values DMSO versus dose. Panel a i) **0.3 nM**: 0.2897; **1 nM**: 3.4·10^-6^; **3 nM**: 3.8·10^-8^; **10 nM**: 7·10^-11^; **30 nM**: 7.3·10^-10^; **100 nM**: 2.4·10^-10^. Panel a ii) **0.3 nM**: 1; **1 nM**: 0.0645; **3 nM**: 0.0136; **10 nM**: 8.2·10^-5^; **30 nM**: 0.0063; **100 nM**: 2.4·10^-6^. (N = 14; 14; 14; 14; 14; 14; 14) Panel a iii) **3 nM**: 0.6533; **10 nM**: 4·10^-5^; **30 nM**: 2·10^-9^; **100 nM**: 1.5·10^-15^. Panel a iv) **3 nM**: 0.00054; **10 nM**: 1.9·10^-11^; **30 nM**: < 2·10^-16^; **100 nM**: < 2·10^-16^. (N = 14; 14; 14; 14; 14) P-values baseline versus dose. Panel b i) **1 nM**: 1; **3 nM**: 1; **10 nM**: 1; **30 nM**: 1; **100 nM**: 1; **300 nM**: 1. Panel b ii) **1 nM**: 1; **3 nM**: 1; **10 nM**: 1; **30 nM**: 1; **100 nM**: 1; **300 nM**: 1. (N = 6; 5; 6; 6; 6; 6; 6) Panel b iii) **3 nM**: 1; **10 nM**: 1; **30 nM**: 1; **100 nM**: 0.00801; **300 nM**: 2.7·10^-9^; **1000 nM**: 1.8·10^-10^. Panel b iv) **3 nM**: 1; **10 nM**: 1; **30 nM**: 1; **100 nM**: 0.00084; **300 nM**: 2.9·10^-11^; **1000 nM**: 1.5·10^-11^. (N = 6; 6; 6; 6; 6; 6; 6) P-values baseline versus dose. Panel c i) **1 nM**: 1; **3 nM**: 1; **10 nM**: 1; **30 nM**: 1; **100 nM**: 1; **300 nM**: 1. Panel c ii) **1 nM**: 1; **3 nM**: 1; **10 nM**: 1; **30 nM**: 1; **100 nM**: 1; **300 nM**: 1. (N = 5; 5; 4; 5; 4; 4; 4) Panel c iii) **3 nM**: 1; **10 nM**: 1; **30 nM**: 1; **100 nM**: 0.00181; **300 nM**: 2.9·10^-6^; **1000 nM**: 1.7·10^-5^.Panel c iv) **3 nM**: 1; **10 nM**: 1; **30 nM**: 1; **100 nM**: 0.54836; **300 nM**: 0.01392; **1000 nM**: 8.2·10^-5^.(N = 5; 5; 4; 5; 5; 5; 3) P-values baseline versus dose. Panel d i) **1 nM**: 1; **3 nM**: 1; **10 nM**: 1; **30 nM**: 0.47; **100 nM**: 1. Panel d ii) **1 nM**: 0.02318; **3 nM**: 0.00170; **10 nM**: 0.00028; **30 nM**: 0.00044; **100 nM**: 0.00113. (N = 5; 5; 5; 5; 5; 5). Panel d iii) **3 nM**: 1; **10 nM**: 1; **30 nM**: 1; **100 nM**: 3·10^-5^. Panel d iv) **3 nM**: 1; **10 nM**: 0.49856; **30 nM**: 0.01473; **100 nM**: 7·10^-6^. (N = 6; 6; 6; 6; 6) P-values Krebs versus dose. Panel e i) **1 nM**: 1; **3 nM**: 1. Panel e ii) **1 nM**: 1; **3 nM**: 0.54. (N = 6; 10; 7) P-values DMSO versus dose. Panel e iii) **10 nM**: 1; **30 nM**: 1; **100 nM**: 1; **300 nM**: 1. Panel e iv) **10 nM**: 0.5298; **30 nM**: 0.2470; **100 nM**: 0.0054; **300 nM**: 0.0029. (N = 7; 8; 4; 5; 7).

Paced (1 Hz) adult rabbit CMs exhibited no significant increase in relaxation time and contraction amplitude at any ISO concentration. At concentrations higher than 3 nM, adult CMs exhibited after-contractions and triggered activity during diastole, which hampered their ability to be paced at a fixed frequency. No significant effects were observed on contraction duration with NIFE, while contraction amplitude significantly decreased in a dose-dependent manner starting from 100 nM (**Fig. 4e**). Data generated by post deflection and sarcomere fractional shortening are available for comparison purposes in **Fig. S6**.

### Analysis of disease phenotypes in vivo

To extend analysis to hearts *in vivo,* we took advantage of the transparency of zebrafish, which allows recording of contracting cardiac tissue *in vivo* (**Fig. 5a, Movie S9**). It was previously shown that mutations in G protein β subunit 5 (*GNB5*) are associated with a multisystem syndrome in human, with severe bradycardia at rest. Zebrafish with loss of function mutations in *gnb5a* and *gnb5b* were generated. Consistent with the syndrome manifestation in patients, zebrafish *gnb5a/gnb5b* double mutant embryos showed severe bradycardia in response to parasympathetic activation^23^. Irregularities in heart rate were visually evident and were clearly distinguishable from the wild type counterpart after analysis with MUSCLEMOTION (**Fig. 5b**). Quantification of the heart rate of these zebrafishes with MUSCLEMOTION highly correlated (R^2^ = 0.98) with the results of the published manual analyses^23^ (**Fig. 5c**). There was however, a striking time-saving for operators in carrying out the analysis using the algorithm (5-10 times faster than manual analysis; 150 recordings were analysed in 5 hours versus 4 days) without compromising accuracy of the outcome. Qualitative analysis of contraction patterns allowed rapid discrimination between arrhythmic vs non-arrhythmic responses to carbachol treatment (**Fig. 5c**).

**Figure 5.**
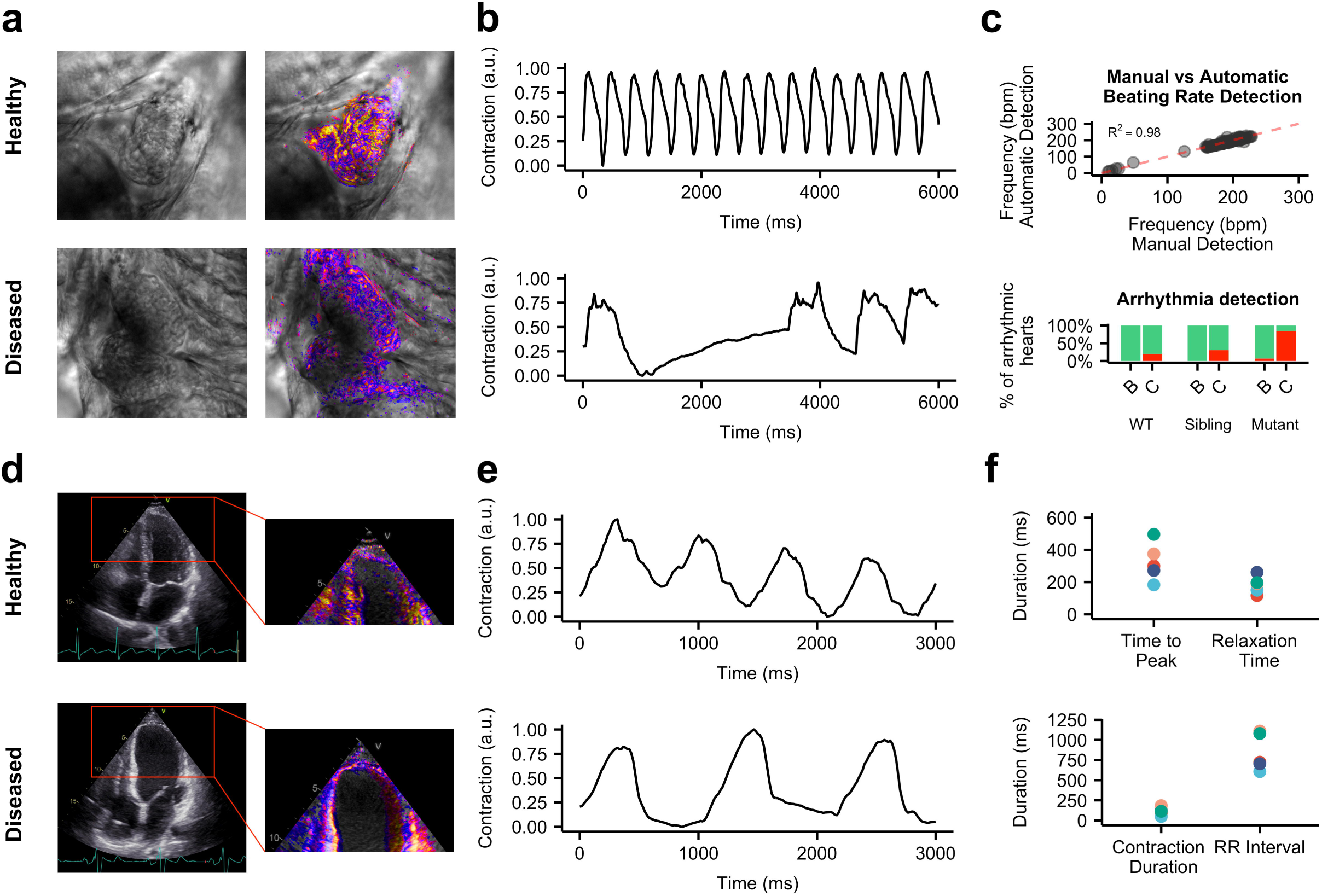
In vivo disease phenotypes. **a)** Representative examples of wild type **(top)** and *gnb5a/gnb5b* mutant **(bottom)** zebrafishes and relative enhancement of moving pixels. **b)** Representative qualitative analyses of normal **(top)** and arrhythmic **(bottom)** contraction profiles from wild type and *gnb5a/gnb5b* mutant zebrafishes treated with carbachol. **c)** Correlation of results obtained from manual (x-axis) vs automatic (y-axis) detection of beating frequency **(top)**; distribution of normal (green) and arrhythmic (red) contraction patterns in baseline condition (B) and after treatment with carbachol (C) in wild type and *gnb5a/gnb5b* mutant zebrafishes **(bottom)**. **d)** Representative echocardiograms of healthy **(top)** and cardiomyopathic **(bottom)** human individuals. Ventricles have been manually cropped and the enhancement of moving pixels is overlaid. **e)** Representative qualitative analyses of normal **(top)** and poor **(bottom)** ventricular functions. **f)** Quantitative data collected from echocardiogram in 5 individuals. Each colour represents one individual.

Finally, we examined human echocardiograms from five healthy and cardiomyopathic individuals (**Fig. 5d**). To assess ventricular function, videos were cropped to exclude movement contributions of the atria and valves. MUSCLEMOTION enabled rapid quantification of temporal parameters from standard ultrasound echography (**Fig. 5e**) such as time-to-peak, relaxation time, RR interval and the contraction duration (**Fig. 5f**).

## Discussion

A reliable and easy-to-use method to quantify cardiac muscle contraction would be of significant benefit to many basic and clinical science laboratories to characterize cardiac disease phenotypes, understand underlying disease mechanisms and predicting cardiotoxic effects of drugs^14^,^24^. Quantification of frame-to-frame differences in pixel intensity has been used in recent reports with success^10^; however, the full spectrum of applications for which these algorithms are relevant, how their output data correlates with gold standards in each system and software performance, specifications, license and software availability, have remained unclear.

Here we developed and tested a user-friendly, inexpensive, open source software platform that serves this purpose in a variety of biological systems of heart tissue. Its integration into current research practices would benefit data sharing, reproducibility, comparison and translation in many clinically relevant contexts^25^.

The linearity and reliability of MUSCLEMOTION were validated using a 3D reconstructed artificial CM which gave the expected linear correlations between known inputs and the outputs (**Fig. 1d-f**). When random repetitive patterns were applied, amplitude outputs differed from inputs, suggesting a potential limitation to measuring contraction amplitudes in highly repetitive biological samples (such as when sarcomere patterns are well-organized), while temporal parameters remained valid (**Fig. 1d,e,g**). However, conditions such as these would be unlikely in standard biological samples, where camera noise significantly reduces the possibility of saturating pixel movement. We partially attenuated this problem by applying, on user demand, a 10-sigma Gaussian blur filter which significantly increased the accuracy of MUSCLEMOTION with highly repetitive structures. Also, to increase reliability, we built in additional controls to detect any mismatches and errors. MUSCLEMOTION can automatically identify and select the reference frame and increase the signal-to-noise-ratio, features which were particularly relevant in reducing user bias and interaction while improving user experience. MUSCLEMOTION is valid in a wide range of illumination conditions without changing temporal parameters; however, exposure time was linearly correlated with contraction amplitude (**Fig. S7**). Batch mode analyses and data storage in custom folders were also incorporated to support overnight automated analyses. For accurate quantification of amplitude, time-to-peak and relaxation time, an appropriate sampling rate should be chosen. For applications similar to those described here, we recommend recording rates higher than 70 frames per second to sample correctly the fast upstroke of the time-to-peak typical of cardiac tissue. This recording rate is easily achievable even using smartphone slow motion video options (∽120/240 frames per second), obviating the need for dedicated cameras and recording equipment if necessary.

We demonstrated excellent linear correlations between our software tool and multiple other standardmethods independent of substrate, cell configuration and technology platform and showed that MUSCLEMOTION is able to capture contraction in a wide range of *in vivo* and *in vitro* applications (**Fig. 2** and **Fig. 3**). Specifically, we identified several advantages compared to optical flow algorithms in terms of speed and the absence of arbitrary binning factors or thresholds which, when modified, profoundly affect the results. One limitation compared to optical flow or EHT standard algorithm is that the tool lacks qualitative vector orientation, making it more difficult to assess contraction direction. Particularly important was the correlation with force data calculated from the displacement of flexible posts by EHTs. This indicates that when the mechanical properties of substrates are known^26^, MUSCLEMOTION allows absolute quantification of contractile force. Technical limitations of the EHT recording system allowed us to analyze only movies with JPEG compression; this resulted in loss of pixel information that might have negatively influenced the correlation shown. For better and more accurate results on contraction quantification, non-lossy/uncompressed video formats should be used for recordings since individual pixel information is lost upon compression and therefore not available for analysis by MUSCLEMOTION.

We proposed and validated practical application in pharmacological challenges using multiple biological preparations recorded in different laboratories; this means that immediate use in multiple independent high-throughput drug-screening pipelines is possible without further software development being required, as recently applied for a drug screening protocol on cardiac organoids from hPSCs^17^. Intuitively, the possibility of having inter-assay comparisons will also be of particular relevance where comparisons of contraction data across multiple platforms are required by regulatory agencies or consortia (e.g. CiPA, CSAHi)^5^,^6^,^22^,^27^. Moreover, this might offer a quantitative approach to investigating how genetic or acquired diseases of the heart (e.g. cardiomyopathies^7^, Long QT Syndrome^28^), heart failure resulting from anticancer treatments^29^,^30^ or maturation strategies^18^,^31^,^32^ affect cardiac contraction. The possibility of linking *in vitro* with *in vivo* assays, with low cost technologies applicable with existing hardware certainly represents an advantage as demonstrated by automatic quantification of zebrafish heartbeats and human echocardiograms (**Fig. 5**). Overall, these results clearly demonstrated that contraction profiles could be derived and quantified in a wide variety of commonly used experimental and clinical settings. MUSCLEMOTION might represent a starting point for a swift screening method to provide clinically relevant insights into regions of limited contractility in the hearts of patients. We encourage further development of this open source platform to fit specific needs; future areas of application could include skeletal or smooth muscle in the same range of formats described here.

MUSCLEMOTION allows the use of a single, transparent method of analysis of cardiac contraction in many modalities for rapid and reliable identification of disease phenotypes, potential cardiotoxic effects in drug screening pipelines and translational comparison of contractile behaviour.

## Limitations

Saturation of pixel movements may affect contraction amplitudes. However, as demonstrated with the artificial CM, contraction velocity and all temporal parameters remained valid. We also minimized the impact of highly repetitive structures on the output of MUSCLEMOTION by applying a Gaussian filter, which also helped in reducing the impact of transverse movements on contraction profiles. High frequency contraction might complicate baseline detection, especially if the duration of the contracted state is similar to that of the relaxed (e.g. approaching sinusoidal). We have implemented a “*fast mode*” option that captures reliable baseline values even at high contraction rates. Furthermore, recordings must be free of moving objects (e.g. debris moved by flow, air bubbles) other than those of interest.

## Acknowledgements

This work was initiated in the context of The National Centre for the Replacement, Refinement and Reduction of Animals in Research (NC3Rs) CRACK IT InPulse project code 35911-259146, with support from GlaxoSmithKline. It was supported by the following grants: ERC-AdG STEMCARDIOVASC (MCL MBJ, GE, BM, TLGJ), ZonMW MKMD *Applications of Innovations 2015-2016* (MCL, BM, SL), BHF SP/15/9/31605 & PG/14/59/31000 and BIRAX 04BX14CDLG grants (DC), ERC-AdG IndivuHeart (ET) and DZHK (German Centre for Cardiovascular Research; ET, SU, HA, MI), ERC-StG StemCardioRisk (DRP, MMPH), VIDI-917.15.303 (the Netherlands Organisation for Scientific Research (NWO); DRP, GC). The Dutch Heart Foundation (CVON 2012 – 10 Predict project), E-Rare (CoHeart project).

## Author Contributions

**SL:** project design, patch clamp, monolayer and organoids experiments, algorithm design, data analysis, statistics, wrote the manuscript.

**MBJ:** project design, monolayer, organoids and membrane labelling experiments, algorithm design, data analysis, statistics, wrote the manuscript.

**TLGJ:** project supervision, algorithm design, optical flow analyses.

**BJ:** supervision of zebrafish experiments.

**BM:** supervision of experiments on isolated hPSC-CM, cardiac organoids and monolayers.

**DRP:** supervision of cell culture for membrane labelling experiments.

**DC:** expert advice coordination of multi-center drug experiments under Crack-IT InPulse.

**DMAE:** designed and rendered the 3D artificial cells.

**ET:** supervision of experiments on Engineered Heart Tissues.

**GE:** generation of cardiac organoids and cell culture.

**CG:** cell culture for membrane labelling experiments.

**HA:** supervision on experiments on engineered heart tissues.

**HER:** advices and supervision on echocardiography data.

**JMRM:** echocardiography recordings and supervision on echocardiography data.

**KSM:** recordings and data analysis of zebrafish hearts.

**KCD:** recordings and data analysis of zebrafish hearts.

**LQ:** recordings and data analysis of adult rabbit cardiomyocytes.

**MI:** experiments and recordings of engineered heart tissues.

**MMPH:** cell culture for membrane labelling experiments.

**OVV:** supervision of experiments on cardiac organoids.

**PR:** supervision of drug tests experiments on aligned cardiomyocytes.

**RMC:** experiments on aligned cardiomyocytes.

**SU:** data analysis of engineered heart tissues.

**SGL:** project supervision and discussion.

**MCL:** project supervision and discussion, wrote the manuscript.

**BFL:** project supervision, algorithm design, discussion.

## Bibliography

Laverty, H. et al. How can we improve our understanding of cardiovascular safety liabilities to develop safer medicines? Br. J. Pharmacol. 163,675–693 (2011).

Passier, R., Orlova, V. & Mummery, C. Complex Tissue and Disease Modeling using hiPSCs. Cell Stem Cell 18,309–321 (2016).

Bellin, M., Marchetto, M. C., Gage, F. H. & Mummery, C. L. Induced pluripotent stem cells: the new patient? Nat Rev Mol Cell Biol 13,726–726 (2012).

van Meer, B. J., Tertoolen, L. G. J. & Mummery, C. L. Concise Review: Measuring Physiological Responses of Human Pluripotent Stem Cell Derived Cardiomyocytes to Drugs and Disease. Stem Cells 34,2015–2015 (2016).

Kitaguchi, T. et al. CSAHi study: Evaluation of multi-electrode array in combination with human iPS cell-derived cardiomyocytes to predict drug-induced QT prolongation and arrhythmia-Effects of 7 reference compounds at 10 facilities. Journal of Pharmacological and Toxicological Methods 78,102–102 (2016).

Hwang, H. S. et al. Comparable calcium handling of human iPSC-derived cardiomyocytes generated by multiple laboratories. J Mol Cell Cardiol 85,88–88 (2015).

Birket, M. J. et al. Contractile Defect Caused by Mutation in MYBPC3 Revealed under Conditions Optimized for Human PSC-Cardiomyocyte Function. Cell Rep 13,745–745 (2015).

Ribeiro, A. J. S. et al. Contractility of single cardiomyocytes differentiated from pluripotent stem cells depends on physiological shape and substrate stiffness. Proc. Natl. Acad. Sci. U.S.A. 112,12710–12710 (2015).

Ribeiro, A. J. et al. Multi-Imaging Method to Assay the Contractile Mechanical Output of Micropatterned Human iPSC-Derived Cardiac Myocytes. Circ Res CIRCRESAHA. 116.310363–91 (2017). doi:10.1161/CIRCRESAHA.116.310363

Kijlstra, J. D. et al. Integrated Analysis of Contractile Kinetics, Force Generation, and Electrical Activity in Single Human Stem Cell-Derived Cardiomyocytes. Stem Cell Reports 5,1238–1238 (2015).

Stoehr, A. et al. Automated analysis of contractile force and Ca2 + transients in engineered heart tissue. Am J Physiol Heart Circ Physiol 306,H1353–H1363 (2014).

Hayakawa, T. et al. Image-based evaluation of contraction–relaxation kinetics of human-induced pluripotent stem cell-derived cardiomyocytes: Correlation and complementarity with extracellular electrophysiology. J Mol Cell Cardiol77,191–191 (2014).

Hayakawa, T. et al. Noninvasive evaluation of contractile behavior of cardiomyocyte monolayers based on motion vector analysis. Tissue Engineering Part C: Methods 118,32–32 (2012).

Mannhardt, I. et al. Human Engineered Heart Tissue: Analysis of Contractile Force. Stem Cell Reports 7,42–42 (2016).

van den Berg, C. W., Elliott, D. A., Braam, S. R., Mummery, C. L. & Davis, R. P. Differentiation of Human Pluripotent Stem Cells to Cardiomyocytes Under Defined Conditions. Methods Mol. Biol. 1353,180–180 (2016).

Sala, L. et al. A new hERG allosteric modulator rescues genetic and drug-induced long-QT syndrome phenotypes in cardiomyocytes from isogenic pairs of patient induced pluripotent stem cells. EMBO Mol Med 8,1081–1081 (2016).

Giacomelli, E. et al. Three-dimensional cardiac microtissues composed of cardiomyocytes and endothelial cells co-differentiated from human pluripotent stem cells. Development dev. 143438 (2017). doi:10.1242/dev.143438

Ribeiro, M. C. et al. Functional maturation of human pluripotent stem cell derived cardiomyocytes in vitro-- correlation between contraction force and electrophysiology. Biomaterials 51,150–150 (2015).

MacQuaide, N., Ramay, H. R., Sobie, E. A. & Smith, G. L. Differential sensitivity of Ca2 + wave and Ca2 + spark events to ruthenium red in isolated permeabilised rabbit cardiomyocytes. Journal of Physiology 588,4742–4742 (2010).

Rocchetti, M. et al. Ranolazine prevents INaL enhancement and blunts myocardial remodelling in a model of pulmonary hypertension. Cardiovascular Research 104,48–48 (2014).

Bers, D. M. Cardiac excitation-contraction coupling. Nature 415,205–205 (2002).

Sala, L., Bellin, M. & Mummery, C. L. Integrating cardiomyocytes from human pluripotent stem cells in safety pharmacology: has the time come? Br. J. Pharmacol. 97,2684 (2016).

Lodder, E. M. et al. GNB5 Mutations Cause an Autosomal-Recessive Multisystem Syndrome with Sinus Bradycardia and Cognitive Disability. Am. J. Hum. Genet. 99,710–710 (2016).

Rodriguez, M. L. et al. Measuring the Contractile Forces of Human Induced Pluripotent Stem Cell-Derived Cardiomyocytes With Arrays of Microposts. J Biomech Eng 136,051010–051010 (2014).

Bullen, A. Microscopic imaging techniques for drug discovery. Nat Rev Drug Discov 7,67–67 (2008).

Vandenburgh, H. et al. Drug-screening platform based on the contractility of tissue-engineered muscle. Muscle Nerve 37,447–447 (2008).

Cavero, I. & Holzgrefe, H. Comprehensive in vitro Proarrhythmia Assay, a novel in vitro/in silico paradigm to detect ventricular proarrhythmic liability: a visionary 21st century initiative. Expert Opin Drug Saf 13,758–758 (2014).

Rocchetti, M. et al. Elucidating arrhythmogenic mechanisms of long-QT syndrome CALM1-F142L mutation in patient-specific induced pluripotent stem cell-derived cardiomyocytes. Cardiovascular Research (2017). doi:10.1093/cvr/cvx006

Burridge, P. W. et al. Human induced pluripotent stem cell-derived cardiomyocytes recapitulate the predilection of breast cancer patients to doxorubicin-induced cardiotoxicity. Nat Med 22,556–556 (2016).

Bellin, M. & Mummery, C. L. Stem cells: The cancer’s gone, but did chemotherapy damage your heart? Nat Rev Cardiol 13, 384–384 (2016).

Nunes, S. S. et al. Biowire: a platform for maturation of human pluripotent stem cell-derived cardiomyocytes. Nat Meth 10,787–787 (2013).

Chan, Y.-C. et al. Electrical stimulation promotes maturation of cardiomyocytes derived from human embryonic stem cells. J Cardiovasc Transl Res 6,999–999 (2013).

